# The impact of sugar diet on humidity preference, survival, and host landing in mosquitoes

**DOI:** 10.1101/2024.09.23.613762

**Authors:** Shyh-Chi Chen, Christopher J. Holmes, Oluwaseun M. Ajayi, Grace Goodhart, Daniel Eaton, Nathan Catlett, Tabitha Cady, Hannah Tran, Luke E. Lutz, Lyn Wang, Ella Girard, Jaida Savino, Amena Bidiwala, Joshua B. Benoit

**Affiliations:** Department of Biological Sciences, University of Cincinnati, Cincinnati, Ohio, USA

**Keywords:** sugar feeding, water seeking, humidity preference, survivorship, mosquito control

## Abstract

Mosquito-borne diseases have caused more than one million deaths each year. There is an urgent need to develop an effective way to reduce mosquito-host interaction to mitigate disease transmission. Sugar diets have long been linked to abnormal physiology in animals, making them potential candidates for mosquito control. Here, we show the impact of sugar diets on humidity preference and survival in *Aedes aegypti* and *Culex pipiens*. With two-choice assays between 100% and 75% relative humidity (RH), we demonstrate that the effect of sugar diets on humidity preference is species-specific where *Ae. aegypti* showed significant differences and the reduced effects were noted in *Cx. pipiens*. Among the sugar diets, arabinose significantly reduced the survival rate of mosquitoes even at low concentrations. Moreover, we found that host landing was not impacted by feeding on different sugar types. Our study suggests that specific sugar treatments could be applied to mosquito control by dampening their humidity preference and reducing their lifespan, thus reducing mosquito-borne disease transmission.

## Introduction

Mosquitoes are the deadliest animals, spreading various mosquito-borne diseases to their hosts (Qureshi 2018; CDC 2023) and claiming many human lives each year (CDC 2023; WHO 2023). Many efforts have been made for mosquito control, and among existing vector control strategies, insecticides play an important role in the intervention in mosquito-borne disease transmission (Qureshi 2018; WHO 2018; CDC 2023; WHO 2023). For example, insecticide applications, such as insecticide-treated mosquito nets (ITNs) and indoor residual spraying (IRS) of insecticides, effectively reduce human mortality rates from malaria (WHO 2018). However, the drawback of insecticide resistance in mosquitoes as well as the potential toxicity to humans and ecosystems have become a growing concern (Hayes et al. 2006; Jayaraj et al. 2016; Moyes et al. 2017; Asgarian et al. 2023). Therefore, there is an urgent need for alternative control approaches for mosquito suppression.

Sugars are a primary dietary source for many insects. For mosquitoes, sugar feeding not only sustains their energy levels but influences many aspects of physiology and behaviors, such as immunity, metabolism, growth, fecundity, longevity, and host-feeding avidity (Edman et al. 1992; Straif and Beier 1996; Gouagna et al. 2014; Phasomkusolsil et al. 2017; Ferguson et al. 2019; Almire et al. 2021; Huck et al. 2021; League et al. 2021). Adult mosquitoes often feed on sugar sources from floral and extrafloral nectaries with preferences for particular types of plants (Yu et al. 2016; Barredo and DeGennaro 2020; Cassone et al. 2024). Most importantly, evidence has shown that sugar feeding in mosquitoes is highly associated with their blood-feeding onsets (Foster 1995). Although sugar diets play an important role in mosquito fitness and survival, evidence in various animal models has shown that high sugar consumption has negative effects on animal health, which can shorten their lifespan (Na et al. 2013; Chandegra et al. 2017; May et al. 2019; van Dam et al. 2020; Catalani et al. 2021; Abe et al. 2022 Oct 13; Pardo-Garcia et al. 2023; Straser et al. 2023; Suzuki et al. 2023). Besides concentration, consumption of varying dietary sugar sources (both natural and artificial) contributes to developmental impacts due to differences in nutritive qualities (Harvey et al. 2012; Li et al. 2020; Morimoto et al. 2020; Force et al. 2023). These studies imply the potential applications of sugar diets for mosquito control when sugar sources are combined with attractive olfactory and visual cues (Scott-Fiorenzano et al. 2017; Dieng et al. 2018).

Attractive toxic sugar baits (ATSBs) are composed of an olfactory attractant, sugar source as a feeding stimulant, and an oral toxicant (Fiorenzano et al. 2017; Kumar et al. 2021; Njoroge et al. 2023). Two advantages make ATSB application a promising strategy for mosquito control: (1) the different configurations of each component result in various types of ATSBs (Stewart et al. 2013; Gu et al. 2020; Kumar et al. 2022; Pullmann-Lindsley et al. 2023) and (2) the combinations can be tailored to specific species, all while avoiding or minimizing toxic effects on non-target species (Fiorenzano et al. 2017; Diarra et al. 2021). After attractants entice animals to land on ATSBs, sugar serves as a feeding stimulant to encourage animals to ingest toxicant, which is often an insecticide at low dosage (Qualls et al. 2014; Barbosa et al. 2019; Kumar et al. 2021). Recent studies indicate that heavy sugar diets or consumption of artificial sweeteners lead to negative impacts on human health and the lifespan or fitness of other animals (Choi et al. 2017; Musso et al. 2017; Gáliková and Klepsatel 2018; Lee et al. 2021; Witek et al. 2022). In *Drosophila*, high sucrose diets are associated with oxidative stress, cardiomyopathy, defective immune responses, dysfunction of intestine homeostasis, obesity, and insulin resistance (Musselman et al. 2011; Na et al. 2013; Zhang et al. 2017; Yu et al. 2018; van Dam et al. 2020; Adesanoye et al. 2021; Catalani et al. 2021). In addition to a high sugar diet, there is a link between the consumption of various sweeteners and an alteration in food intake, sleep, memory, learning, and neural development in *Drosophila* (Wang et al. 2016; Hasegawa et al. 2017; Musso et al. 2017; Park et al. 2017; Santos-Cruz et al. 2023). Most importantly, some of these sugar substitutes possess insecticidal properties that significantly reduce mosquito survivorship (Sharma et al. 2020; Maestas et al. 2023; Pullmann-Lindsley et al. 2023); therefore, high-sugar diets and artificial sweeteners provide potential value in mosquito control by serving a hybrid role as both a stimulant and toxicant in ATSBs. However, outside of the impact on survival, little is known about whether these sugar and sweetener diets impact other areas of mosquito biology and the variation between species. In the present study, we investigated how various sugar diets affect the humidity preference, blood feeding, and survivorship of the yellow fever mosquito, *Ae. aegypti,* and the northern house mosquito, *Cx. pipiens*. We found that these treatments impair mosquito humidity preference and survivorship in a species-specific manner, but do not impact host landing by mosquitoes. Our understanding of the effects of sugar diet treatments may help us develop better strategies for the application of ATSBs in targeted mosquito species. This research fulfills urgent needs for novel strategies that do not contribute to insecticide resistance, to reduce mosquito population and mitigate the burden of mosquito-borne diseases.

## Materials and Methods

### Mosquito husbandry and diet preparation

*Ae. aegypti* (Gainesville) and *Cx. pipiens* (Buckeye) were kept in the insectary with 15h:9h light:dark cycles and at constant temperature (25°C) and humidity (∼80%RH). Mosquito larvae were reared with fish food (Tetramin) mixed with yeast extract. Adult mosquitoes were kept in the 30 x 30 x 30 cm cages and provided with deionized water and 10% sucrose solution *ad libitum* before sugar treatment (sucrose, D- and L-arabinose: Thermo Fisher Scientific; D-sorbitol: Acros Organics; sucralose: Midwood Brands). When mosquitoes were 10-12-day-old, the deionized water and 10% sucrose solution were replaced with various sugar diets (Supp. Table 1 and 2) for 5-day sugar diet treatment. Sugar treatment includes the control (10% sucrose + water) and various sugar diet groups (30% sucrose + water, 10% sucrose only, 30% sucrose only, 10% arabinose + water, 10% sucralose + water, 10% sorbitol only, 10% sorbitol + water).

### Avidity and behavioral preference

Following the 5-day sugar treatments, ∼25-35 *Ae. aegypti* and *Cx. pipiens* female mosquitoes were transferred to a 30 x 30 x 30 cm cage for two-choice behavioral assays (see Fig. 1 for the setup). This behavioral cage includes two jars with deionized water and oversaturated NaCl solution for 100% and 70% humidities, respectively. Twenty-four hours after two-choice behavioral assays, the humidity avidity and behavioral preference index (PI) of mosquitoes were calculated as follows:

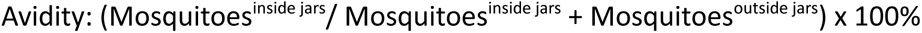

Behavioral preference index (PI): Mosquito Numbers (100% - 75% RH)/Mosquito Numbers (100% + 75% RH)

**Fig. 1.**
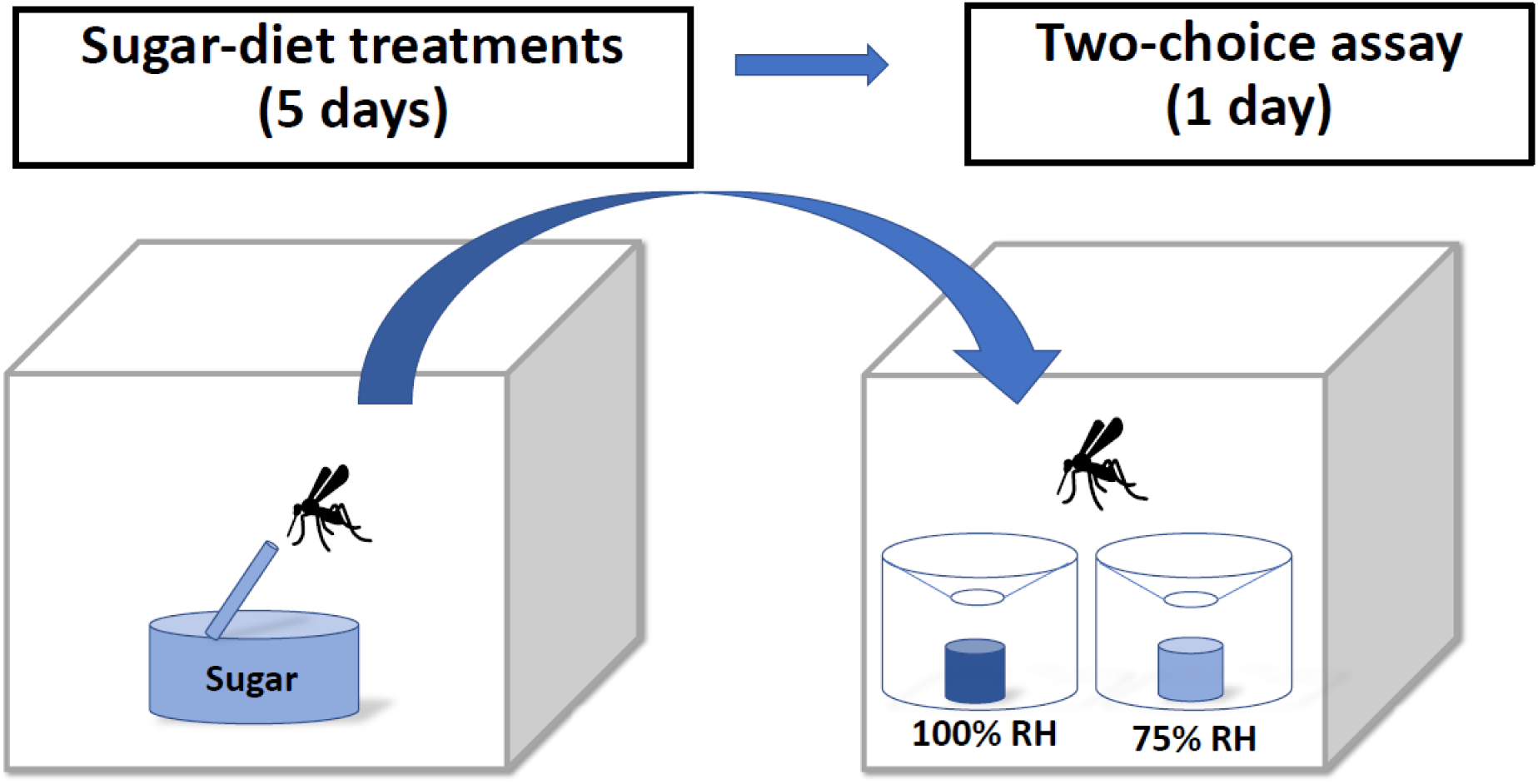
The experimental design for sugar-diet treatments and behavioral assays. After feeding on sugar diets for 5 days, mosquitoes were transferred to a cage for two-choice assays. The cage contained two jars with high (100%) and low (70%) humidities.

### Survival rate of *Ae. aegypti* and *Cx. pipiens* in different concentrations of arabinose

To assess the effect of sugar diet on mosquito survivorship, 10-20 female mosquitoes (10-12-day-old) in the mesh 15 x 15 x 15 cm cages (Bugdorm) were fed with D- and L-arabinose at various concentrations. The number of dead mosquitoes was counted daily until all mosquitoes died in the cage. Generalized additive models (GAMs) were created with the MGCV package (Wood 2004; Wood et al. 2016; Wood 2017) and compared with Tukey’s honestly significant difference pairwise via the emmeans package (Lenth et al. 2024) in R. The ecotox package (Hlina et al. 2019) was used to establish the LT25, LT50, LT75, and LT99 for each GAM in R. Data processing was completed in R (R Core Team 2022) with plyr (Wickham 2011), tidyr (Wickham et al. 2024), dplyr (Wickham et al. 2023), and Rmisc (Hope, M. Ryan 2013). Survival figures were generated in R with ggplot2 (Wickham 2009), specific comparisons with accompanying statistics are included in Supp. Figs. 3-8, and all survival comparison statistics are included in supplemental Tables 3-8).

### Host-seeking behavioral assays for *Ae. aegypti*

Among sugar diets, 30% sucrose, 10% sucralose + water, and 10% sorbitol show significant differences from the control group (10% sucrose + water). To estimate the effect of these sugar diets on host-seeking behaviors, 10 adult *Ae. aegypti* females (10-12-day-old) in 15 x 15 x 15 cm cages were treated with sugar diets for 5 days and then the host-seeking behavioral assays were carried out between 2-5 hours after lights-on (zeitgeber time, ZT2-5). The hand of a volunteer was used to provide thermal and olfactory cues, and a 1-cm circular spacer was placed between the hand and the cage to prevent direct host contact with mosquitoes (Laursen et al. 2023). Human trials are approved by the University of Cincinnati (IRB 2021-0971, University of Cincinnati). During the entire 5-minute assay period, the number of mosquitoes landing on the spacer area below hand was counted as follows:

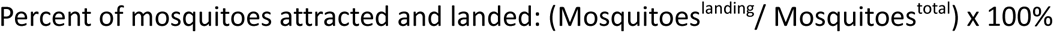

## Results

### Avidity-fitness and behavioral preference index (PI)

To determine how sugar diets impact mosquito humidity preference, we used two-choice assays and assessed the water-seeking behaviors of *Ae. aegypti* and *Cx. Pipiens* (Hayes et al. 2006; Moyes et al. 2017; Asgarian et al. 2023). We found that *Ae. aegypti* and *Cx. pipiens* showed different profiles of humidity avidity after 5-day sugar treatments. Compared to the control group (10% sucrose + water), all sugar diets but 10% sorbitol + water significantly decreased the humidity avidity of *Ae. aegypti* (i.e. less mosquitoes entered any of the two jars with 70% and 100% RH; Figs. 1 and 2A). *Cx. pipiens,* feeding on 30% sucrose, 10% sucralose + water, and 10% sorbitol + water showed decreased avidity compared to the control group (Fig. 2B). Notably, none of the *Cx. pipiens* treated with 10% D-arabinose + water survived for up to 5 days after the treatment. These results suggest that various sugar diets impact the ability to locate or preference for water sources.

**Fig. 2.**
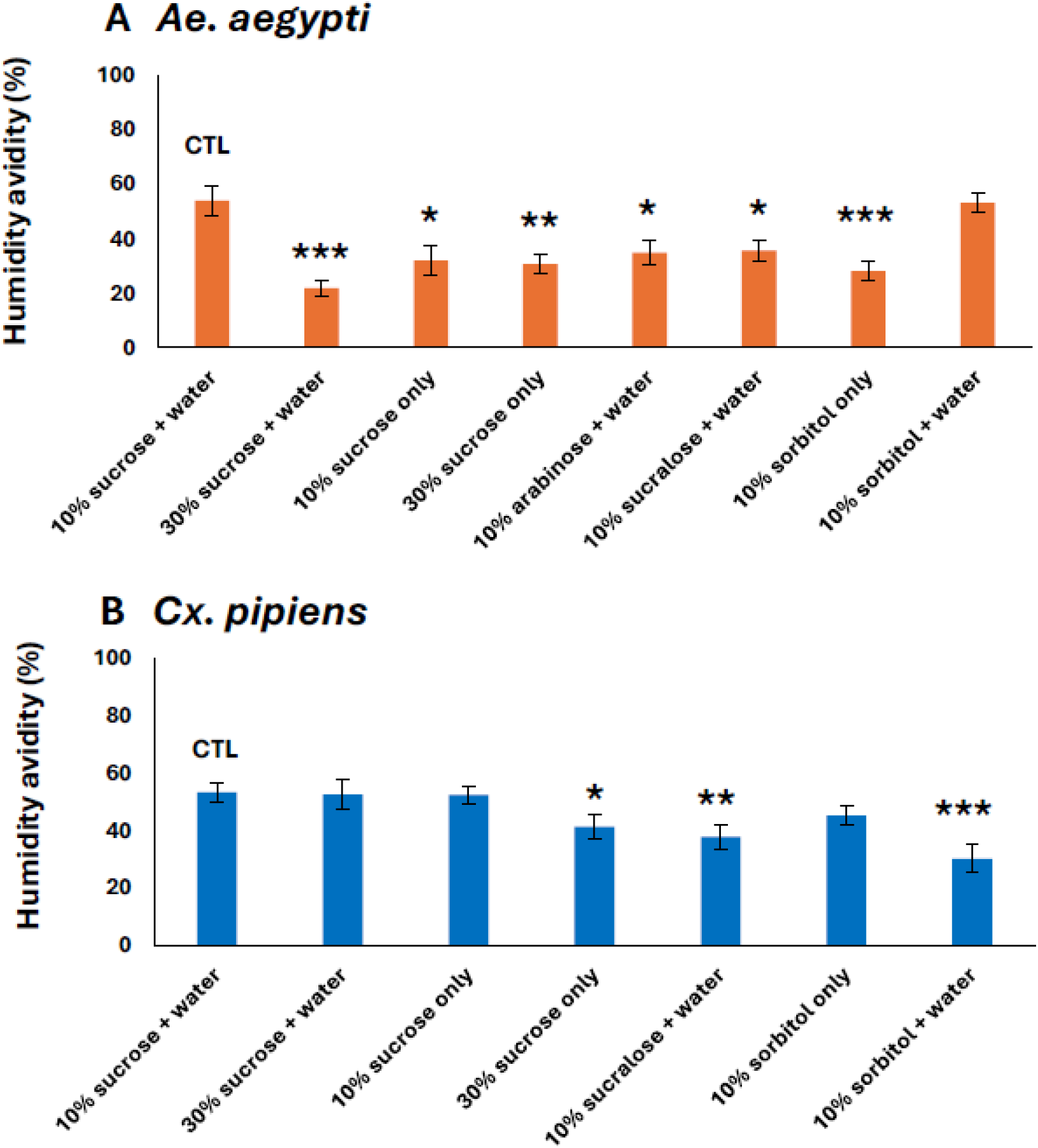
Effect of sugar diets on humidity avidity in mosquitoes after five days of sugar-diet treatments. (A) Compared to the control group (10% sucrose + water), the avidity of *Ae. aegypti* was significantly lower under all but the sorbitol treatment (10% sorbitol + water) (Suppl Table 1). (B) The avidity of *Cx. pipiens* was merely significantly lower in three treatments (30% sucrose only, 10% sucralose + water, and 10% sorbitol + water) (Suppl Table 1). No *Cx. pipiens* survived five days after feeding on 10% arabinose. Statistical difference (t-test) from the control: *, P < 0.05; **, P < 0.01 ***, P < 0.001. For sample size, see Suppl Table 1.

Following the avidity to water sources was calculated, we investigated whether mosquito humidity preference is influenced by sugar diets by comparing how mosquitoes choose high humidity (100%) over low humidity (75%) between the control and sugar treatment groups. The behavioral preference index of *Ae. aegypti* fed on a high sugar diet (30% sucrose) and sweeteners (10% D-arabinose + water, 10% sucralose + water, and 10% sorbitol) were significantly lower than the control group (Fig. 3A). On the contrary, there was no significant difference of behavioral preference index in *Cx. pipiens* with various sugar treatments (Fig. 3B). The results indicate that sugar diets may selectively disturb humidity differentiation in *Ae. aegypti*.

**Fig. 3.**
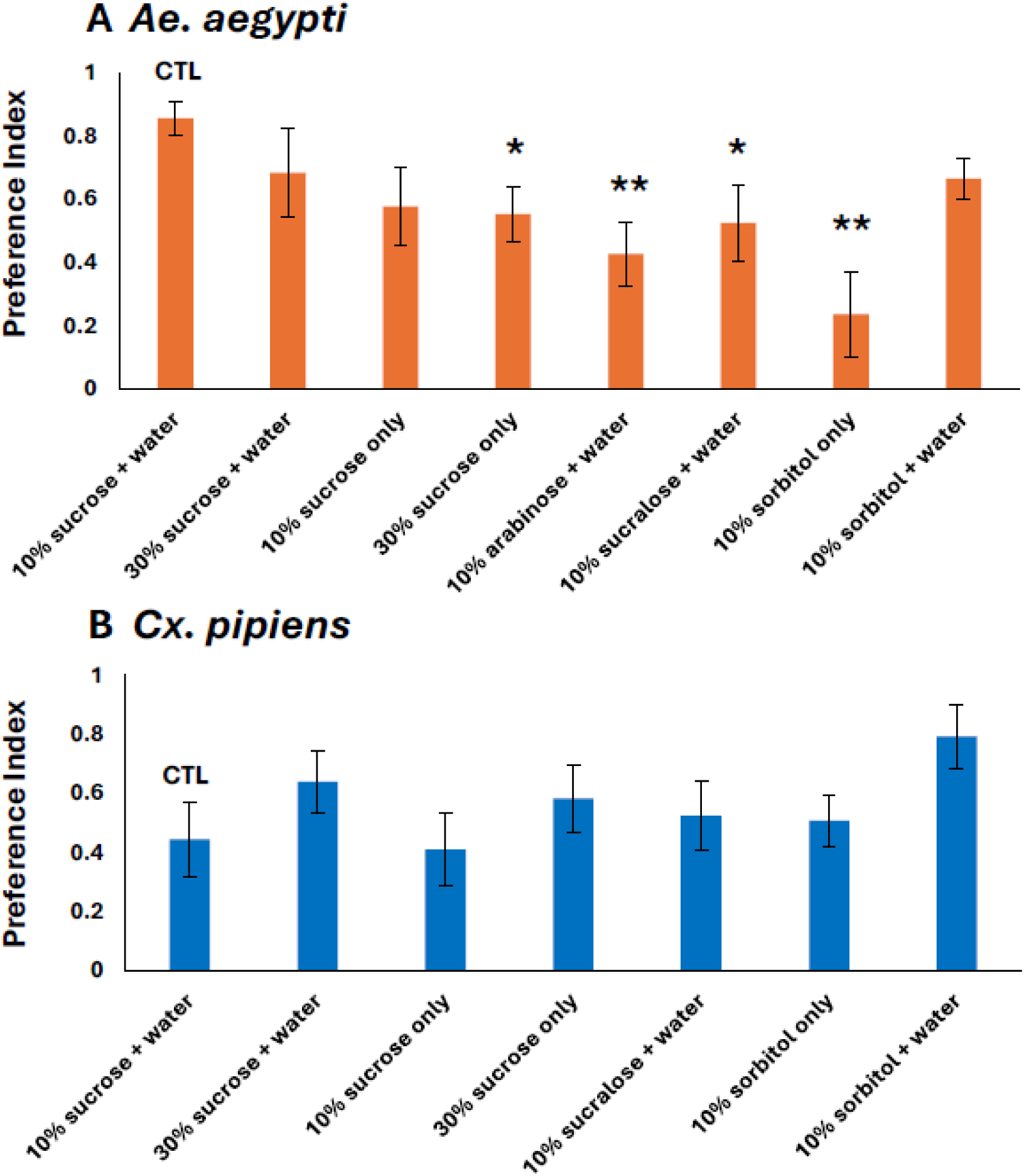
Mosquito behavior preference for humidity after five days of sugar-diet treatments. Preference index (PI) for *Ae. aegypti* (A) and *Cx. Pipiens* (B) was measured from the mosquitoes entering containers with 100%RH in Fig. 2 (for calculations, please see Materials and Methods). (A) *Ae. aegypti* under 30% sucrose only, 10% arabinose + water, 10% sucralose + water, and 10% sorbitol only treatments showed significantly lower PI (Suppl Table 2). (B) *Cx. pipiens* PI had no significant difference under any sugar treatment (Suppl Table 2). *Cx. pipiens* fed on 10% arabinose did not survive five days post-treatment. Statistical difference (t-test) from the control: *, P < 0.05; **, P < 0.01. For sample size, see Suppl Table 2.

### Survival rate for arabinose (LD50)-lifespan/survivorship

Based on our initial observation, high mortality was observed for arabinose during the studies on humidity preference, which we examined in more detail in subsequent studies. Low doses of arabinose appear to decrease estimated GAM survival from the non-arabinose controls in both *Ae. aegypti* and *Cx. pipiens* (Fig. 4 A, C; Supp. Tables 3-4). Estimated mean survival times (LTs) for both *Ae. aegypti* and *Cx. pipiens* were significantly earlier in all arabinose groups when compared to 10% sucralose + water (control) (Fig. 4 B, D; Supp. Tables 5-8).

**Fig. 4.**
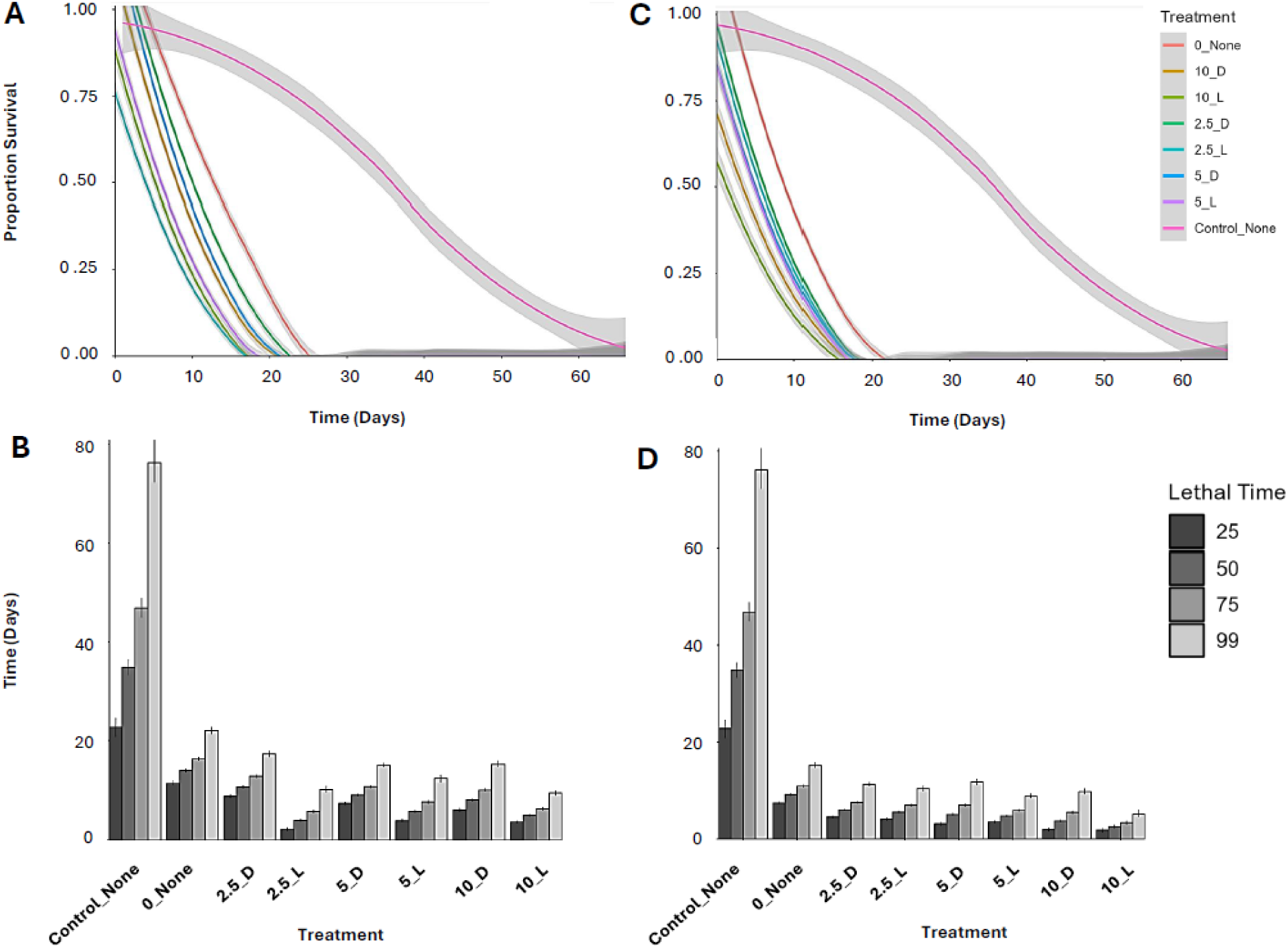
Survival assays for mosquitoes with various arabinose treatments. Generalized additive models (GAMs) and lethal time estimates (LTs) for survival on various arabinose treatments in (A, B) *Ae. aegypti* and (C, D) *Cx. pipiens*. (A, C) GAM arabinose treatments are distinguished by line color and variation estimates are visualized by shading. All significant GAM comparisons are included in Supp. Tables 3-4. (B, D) LT estimates corresponding to LTs 25, 50, 75, and 99 for various arabinose treatments. Median LTs are included in Supp. Tables 5-6 and significantly different LT comparisons of various arabinose treatments are included in Supp. Tables 7-8. Treatment groups: Control_None (10% sucrose + water), 0_None (water), 2.5_D (2.5% D-arabinose + water), 2.5_L (2.5% L-arabinose + water), 5_D (5% D-arabinose + water), 5_L (5% L-arabinose + water), 10_D (10% D-arabinose + water), and 10_L (10% L-arabinose + water).

### Host-seeking behaviors

Our two-choice assay showed that the sugar diets differentially impact the water-seeking behaviors of two mosquito species (Figs. 2 and 3). Humidity detection has been previously shown to impact host seeking in mosquitoes. The results from host attraction assays showed no difference between the control group and mosquitoes feeding on high-sugar diets or sweeteners (Fig. 5, Supp. Table 9).

**Fig. 5.**
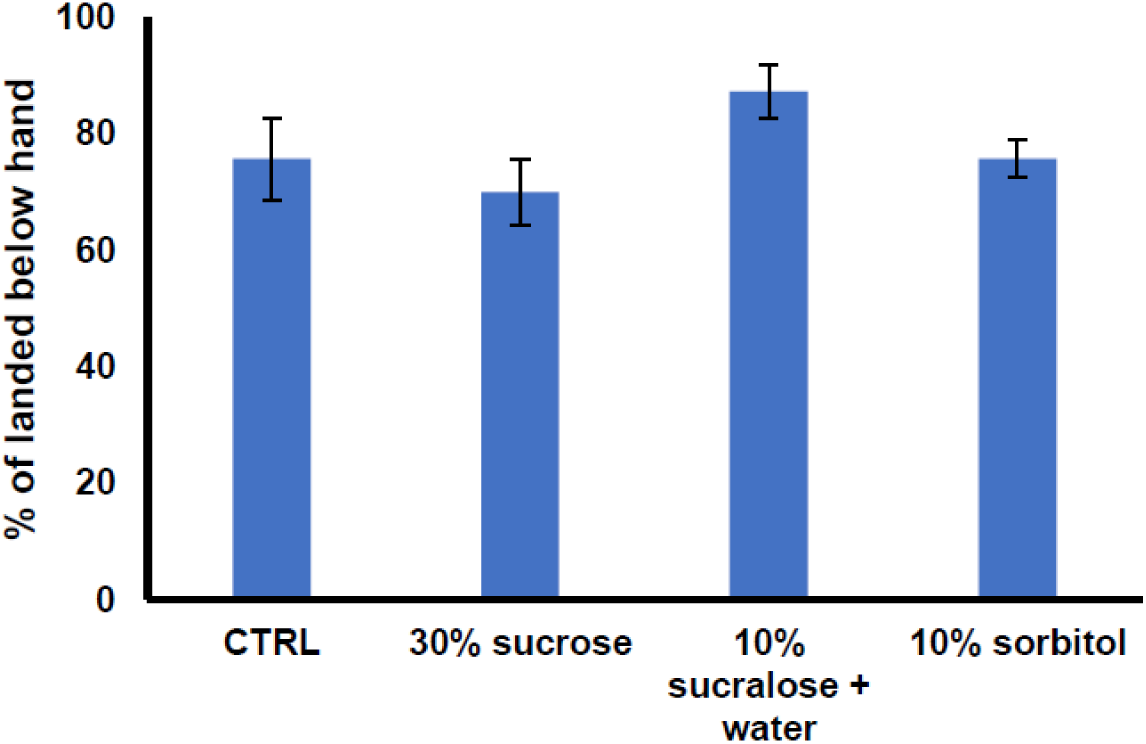
Host-seeking behaviors of *Ae. aegypti* after five days of sugar-diet treatments. The percentage of mosquitoes landing on the spacer area below a host’s hand within the 5-minute behavioral assay. Replicates (n=10 for each treatment), CTRL=7, 30% sucrose= 12, 10% sucralose + water=11, 10% sorbitol=7. T test results show no significance between the control group (CTRL, 30% sucrose + water) and each treatment group (Suppl Table 9).

## Discussion

Sugar is an important carbohydrate source for mosquito energy reserves (Barredo and DeGennaro 2020; Santos et al. 2024). Various types of sugar have been incorporated into ATSBs as sugar baits to stimulate the consumption of oral toxicants and enhance the toxic effects. In combination with miscellaneous attractants and active ingredients, the application of ATSBs can target specific insects and avoid non-target species. Moreover, since some sugar diets are known to be harmful to an animal’s fitness and survival, they can play dual roles of feeding stimulant and toxicant in ATSBs. The advantage of this application is that it reduces the use of toxic pesticides and further prevents pesticide resistance. So far, most research has studied the effects on the target and non-target insects; however, the mechanisms remain largely unknown.

Excess sugar or artificial sweeteners may result in the alteration of insect sensory physiology. For example, high sugar diet reshapes the sweet responses and temperature preference in *Drosophila* within 2-3 days (May et al. 2019; Vaziri et al. 2020). Insects sense sweetness through a conserved family of inotropic gustatory receptors (Grs) which can respond to specific tastants (Freeman et al. 2014; Gomes et al. 2024). The molecular basis of sugar selectivity in Grs is critical to metabolic surveillance and sugar absorption facilitation because not all sugars provide equivalent palatability, nutritive and metabolic values for insects (Burke and Waddell 2011; Fujita and Tanimura 2011). Although specific sugars selectively activate Grs, there is interaction between sugar substances through binding to the same receptors. For example, silkworm BmGr-9 can be selectively activated by D-fructose and other hexoses (glucose, galactose, and sorbose) can inhibit the fructose-activated BmGr-9 activity through non-selective binding (Sato et al. 2011; Gomes et al. 2024). Besides, the sensation of taste in sugars may differ in animals. Many non-caloric sweeteners to humans are much sweeter than sucrose (Świąder et al. 2009; Chattopadhyay et al. 2014) and insects show consumption preference toward specific types of sweeteners depending on the species (King et al. 2020) and this characteristic of sweeteners can be broadly applied to ATSBs for various pest species. Moreover, evidence shows that sweeteners cause dehydration, altered gut microbiome, and osmotic stress in German cockroach and spotted-wing drosophila (Price et al. 2022; Lee et al. 2024). The present study focuses on how sugar diets impact humidity preference in two mosquito species as mosquitoes heavily rely on water detection in various aspects of their life cycles (Brown et al. 2023; Laursen et al. 2023; Verhulst and Mathis 2023). We found that mosquito water-seeking behaviors are differentially affected by sugar diets including high sugar diets and sweeteners. Our results provide additional evidence of sweetener-induced behavioral and physiological disturbance and further support the concept of sweeteners as potential safe insecticides.

Although arabinose offers no nutritive value (Nayar and Sauerman 1971), choice assays showed similar *Ae. aegypti* feeding rates on arabinose-based toxic sugar baits, and blood feeding did not mitigate the mortality of the arabinose-fed mosquitoes (Airs et al. 2019). In choice assays, although sucrose was preferred over arabinose in *Ae. aegypti* (Airs et al. 2019), *Anopheles quadrimaculatus* and *Aedes albopictus* females preferred arabinose, maltose, meliniose, and trehalose over other sugars such as fructose, glucose, melezitose, raffinose, rhamnose, sucrose, and turanose (Xue et al. 2022). Unfortunately, toxicity to arabinose was also noted in non-mosquito species such as the western honeybee, *Apis mellifera* (Barker and Lehner 1974), the boll weevil, *Anthonomus grandis* (Nettles 1972), and the sweet potato whitefly, *Bemisia tabaci* (Hu et al. 2010). However, reduced concentrations of arabinose or the addition of sucrose to the mixture mitigated the mortality levels of these species to some degree (Nettles 1972; Barker and Lehner 1974; Hu et al. 2010). Also, to enhance the attraction of the arabinose-based toxic sugar baits to mosquitoes, additional attractants (CO_2_) or odor cues, such as specific compounds from plant headspace volatiles, could be used based on the preference of mosquito species (Jerry et al. 2017; Barredo and DeGennaro 2020; Madang et al. 2022). Our results show that although 2.5% L-arabinose had a lower LT50 than the 2.5% D-arabinose treatment, there were no differences between different concentrations of L-arabinose in *Ae. aegypti* (Fig. 4, Supp. Table 7), indicating that mortality associated with the L-enantiomer of arabinose was not concentration-dependent at the levels we investigated. No differences were found in *Cx. pipiens* between enantiomers nor concentrations, although 10% L-arabinose trended towards reduced survival (Fig. 4). Therefore, combining 2.5% L-arabinose with complementary additives for toxic bait traps could be particularly effective against mosquitoes.

Many terrestrial insects experience continuous water loss from their body and must find water sources to avoid mortality due to dehydration. Under dry or dehydration conditions, mosquitoes show molecular and behavioral alterations for water loss reduction and rehydration (Benoit et al. 2010; Hagan et al. 2018; Holmes et al. 2022; Holmes et al. 2023). Using sugars with insecticidal activity, such as arabinose, mannose, and xylose, that display antifeedant effects (Hu et al. 2010) in ATSBs may be particularly useful for mosquito control during dry conditions. The present study shows that various sugar diets lead to the species-specific interference of mosquito humidity preference and survival. In combination with other chemical or physical attractants, sugar diets show a potential role in the development of effective species-specific ATSBs, which can be less detrimental to non-targeted animals and the environment than non-specific insecticidal treatments. As insecticide resistance threatens ongoing mosquito control efforts, continued examination of alternative methods, such as ATSBs, is necessary.

## Acknowledgments

We thank the staffs at the University of Cincinnati for their assistance in developing the mesocosm and devices used in this research. Research reported in this publication was supported by the National Institute of Allergy and Infectious Diseases under Award Numbers R01AI148551, R21AI166633, and R21AI176098 (JBB).

## Author contributions

Conceptualization, Supervision: S.C.C. and J.B.B. Visualization, Writing, and Formal analysis: S.C.C. and C.J.H. Funding acquisition: J.B.B. Investigation: O.M.A., G.C., D.E., N.C., T.C., H.T., L.E.L., L.W., E.G., J.S., A.B.

## Supplementary Material

**Suppl. Table 1:**
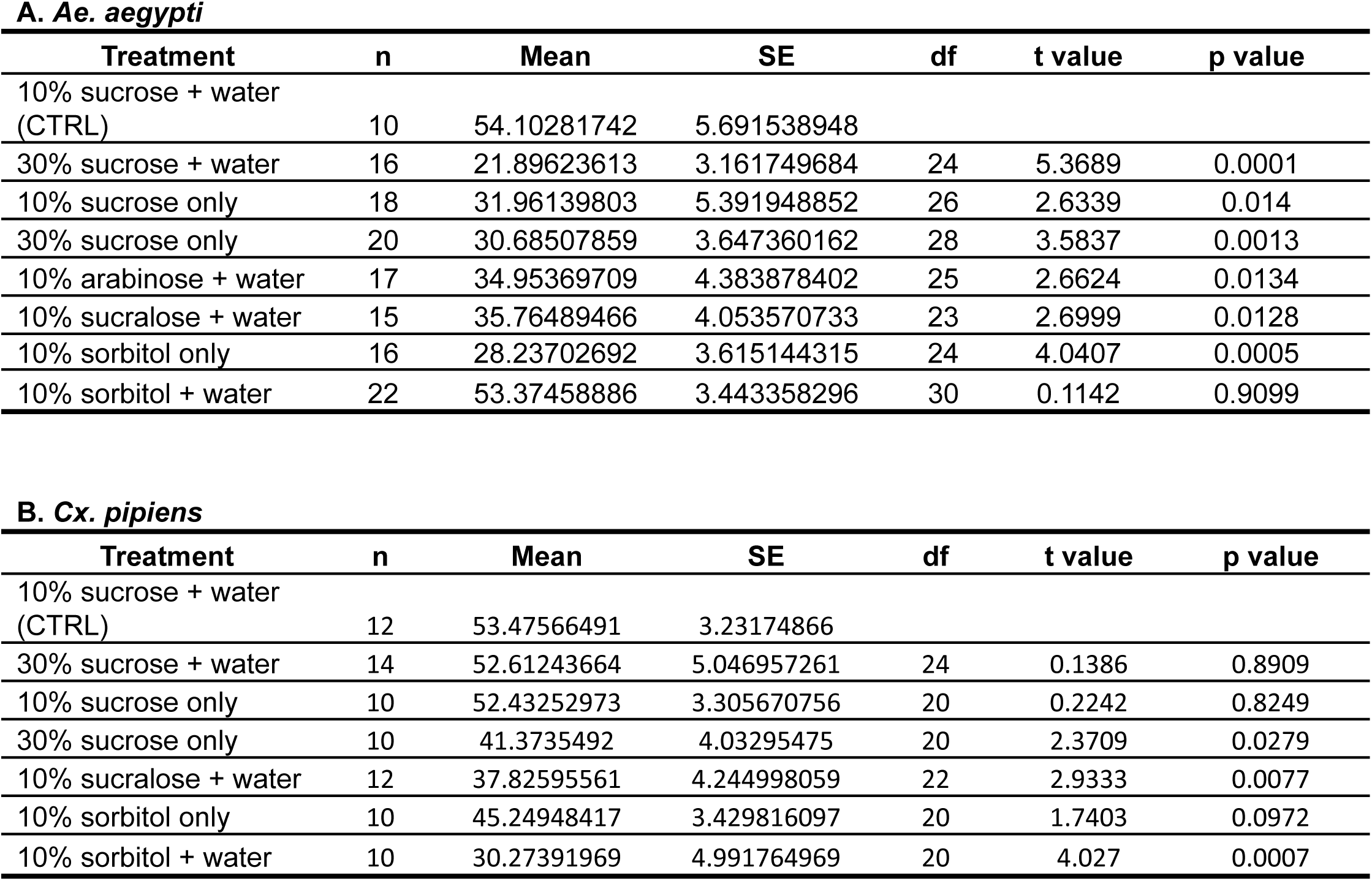
Effect of sugar diets on humidity avidity in mosquitoes after five days of sugar-diet treatments. Compared to the control group (10% sucrose + water), the avidity of (A) *Ae. aegypti* and (B) *Cx. Pipiens* with various sugar diet treatments by t-test.

A. *Ae. aegypti*
| Treatment | n | Mean | SE | df | t value | p value |
| --- | --- | --- | --- | --- | --- | --- |
| 10% sucrose + water (CTRL) | 10 | 54.10281742 | 5.691538948 |  |  |  |
| 30% sucrose + water | 16 | 21.89623613 | 3.161749684 | 24 | 5.3689 | 0.0001 |
| 10% sucrose only | 18 | 31.96139803 | 5.391948852 | 26 | 2.6339 | 0.014 |
| 30% sucrose only | 20 | 30.68507859 | 3.647360162 | 28 | 3.5837 | 0.0013 |
| 10% arabinose + water | 17 | 34.95369709 | 4.383878402 | 25 | 2.6624 | 0.0134 |
| 10% sucralose + water | 15 | 35.76489466 | 4.053570733 | 23 | 2.6999 | 0.0128 |
| 10% sorbitol only | 16 | 28.23702692 | 3.615144315 | 24 | 4.0407 | 0.0005 |
| 10% sorbitol + water | 22 | 53.37458886 | 3.443358296 | 30 | 0.1142 | 0.9099 |

B. *Cx. pipiens*
| Treatment | n | Mean | SE | df | t value | p value |
| --- | --- | --- | --- | --- | --- | --- |
| 10% sucrose + water (CTRL) | 12 | 53.47566491 | 3.23174866 |  |  |  |
| 30% sucrose + water | 14 | 52.61243664 | 5.046957261 | 24 | 0.1386 | 0.8909 |
| 10% sucrose only | 10 | 52.43252973 | 3.305670756 | 20 | 0.2242 | 0.8249 |
| 30% sucrose only | 10 | 41.3735492 | 4.03295475 | 20 | 2.3709 | 0.0279 |
| 10% sucralose + water | 12 | 37.82595561 | 4.244998059 | 22 | 2.9333 | 0.0077 |
| 10% sorbitol only | 10 | 45.24948417 | 3.429816097 | 20 | 1.7403 | 0.0972 |
| 10% sorbitol + water | 10 | 30.27391969 | 4.991764969 | 20 | 4.027 | 0.0007 |

**Suppl. Table 2:** Mosquito behavior preference for humidity after five days of sugar-diet treatments. Compared to the control group (10% sucrose + water), the preference index (PI) of (A) *Ae. aegypti* and (B) *Cx. Pipiens* with various sugar diet treatments by t-test.

A. *Ae. aegypti*
| Treatment | n | Mean | SE | df | t value | p value |
| --- | --- | --- | --- | --- | --- | --- |
| 10% sucrose + water (CTRL) | 10 | 0.855665 | 0.052205 |  |  |  |
| 30% sucrose + water | 15 | 0.682593 | 0.140726 | 23 | 0.9688 | 0.3427 |
| 10% sucrose only | 17 | 0.576946 | 0.122259 | 25 | 1.6842 | 0.1046 |
| 30% sucrose only | 20 | 0.553215 | 0.088895 | 28 | 2.2928 | 0.0296 |
| 10% arabinose + water | 17 | 0.426758 | 0.102394 | 25 | 3.0577 | 0.0053 |
| 10% sucralose + water | 15 | 0.52483 | 0.120831 | 23 | 2.1357 | 0.0436 |
| 10% sorbitol only | 16 | 0.236937 | 0.136272 | 24 | 3.4676 | 0.002 |
| 10% sorbitol + water | 22 | 0.665775 | 0.066174 | 30 | 1.8107 | 0.0802 |

B. *Cx. pipiens*
| Treatment | n | Mean | SE | df | t value | p value |
| --- | --- | --- | --- | --- | --- | --- |
| 10% sucrose + water (CTRL) | 12 | 0.444011 | 0.12774 |  |  |  |
| 30% sucrose + water | 14 | 0.638605 | 0.104648 | 24 | 1.19 | 0.2457 |
| 10% sucrose only | 10 | 0.410594 | 0.120483 | 20 | 0.1876 | 0.8531 |
| 30% sucrose only | 10 | 0.581627 | 0.114434 | 20 | 0.7874 | 0.4403 |
| 10% sucralose + water | 12 | 0.523187 | 0.116092 | 22 | 0.4587 | 0.651 |
| 10% sorbitol only | 10 | 0.507325 | 0.086996 | 20 | 0.3927 | 0.6987 |
| 10% sorbitol + water | 10 | 0.792273 | 0.106454 | 20 | 2.0418 | 0.0546 |

**Supp. Table 3:** Generalized additive model (GAM) comparisons for survival with various arabinose treatments in *Ae. aegypti*.

| Contrast | Estimate | SE | t-ratio | p-value |
| --- | --- | --- | --- | --- |
| 5_D - Control_None | 0.028179 | 0.006703 | 4.204005 | 0.000708 |
| 5_L - Control_None | 0.022544 | 0.00685 | 3.291236 | 0.022501 |
| 2.5_D - Control_None | 0.024936 | 0.007719 | 3.23027 | 0.027374 |
| 10_D - Control_None | 0.026345 | 0.008484 | 3.105338 | 0.040312 |
| 10_L - Control_None | 0.023003 | 0.007768 | 2.961115 | 0.061458 |
| 2.5_L - Control_None | 0.021748 | 0.007763 | 2.801608 | 0.094862 |
| 0_None - Control_None | 0.018421 | 0.008054 | 2.287074 | 0.300934 |

Treatment groups: Control\_None (10% sucrose + water), 0\_None (water), 2.5\_D (2.5% D-arabinose + water), 2.5\_L (2.5% L-arabinose + water), 5\_D (5% D-arabinose + water), 5\_L (5% L-arabinose + water), 10\_D (10% D-arabinose + water), and 10\_L (10% L-arabinose + water).

**Supp. Table 4:** Generalized additive model (GAM) comparisons for survival with various arabinose treatments in *Cx. pipiens*.

| Contrast | Estimate | SE | t-ratio | p-value |
| --- | --- | --- | --- | --- |
| **2.5_D - Control_None** | 0.023653 | 0.005532 | 4.275848 | 0.000518 |
| **0_None - Control_None** | 0.027239 | 0.007847 | 3.471158 | 0.012284 |
| **5_D - Control_None** | 0.022294 | 0.006477 | 3.44177 | 0.013597 |
| **2.5_L - Control_None** | 0.023128 | 0.00714 | 3.239286 | 0.026599 |
| **5_L - Control_None** | 0.022874 | 0.007253 | 3.153728 | 0.034781 |
| **10_D - Control_None** | 0.021366 | 0.008024 | 2.662749 | 0.13452 |
| **10_L - Control_None** | 0.018649 | 0.008375 | 2.226724 | 0.335789 |

Treatment groups: Control\_None (10% sucrose + water), 0\_None (water), 2.5\_D (2.5% D-arabinose + water), 2.5\_L (2.5% L-arabinose + water), 5\_D (5% D-arabinose + water), 5\_L (5% L-arabinose + water), 10\_D (10% D-arabinose + water), and 10\_L (10% L-arabinose + water).

**Supp. Table 5:** Median lethal times (LT50) for various arabinose treatments in *Ae. aegypti* including the lower control limit (LCL), upper control limit (UCL), and standard error (SE) in days.

| Treatment | LT50 | LCL | UCL | SE |
| --- | --- | --- | --- | --- |
| **Control_None** | 34.76196 | 33.25696 | 36.27553 | 0.466949 |
| **0_None** | 13.92029 | 13.63625 | 14.20432 | 0.144523 |
| **2.5_D** | 10.85108 | 10.6138 | 11.08833 | 0.107802 |
| **2.5_L** | 4.108325 | 3.863436 | 4.34203 | 0.121522 |
| **5_D** | 9.208946 | 9.04666 | 9.371202 | 0.082647 |
| **5_L** | 5.912006 | 5.683385 | 6.137363 | 0.115437 |
| **10_D** | 8.211862 | 7.965404 | 8.457422 | 0.125153 |
| **10_L** | 5.084023 | 4.899229 | 5.267589 | 0.093594 |

Treatment groups: Control\_None (10% sucrose + water), 0\_None (water), 2.5\_D (2.5% D-arabinose + water), 2.5\_L (2.5% L-arabinose + water), 5\_D (5% D-arabinose + water), 5\_L (5% L-arabinose + water), 10\_D (10% D-arabinose + water), and 10\_L (10% L-arabinose + water).

**Supp. Table 6:** Median lethal times (LT50) for various arabinose treatments in *Cx. pipiens* including the lower control limit (LCL), upper control limit (UCL), and standard error (SE) in days.

| **Treatment** | **LT50** | **LCL** | **UCL** | **SE** |
| --- | --- | --- | --- | --- |
| **Control_None** | 34.80398 | 33.33385 | 36.2825 | 0.465637 |
| **0_None** | 9.156567 | 8.948662 | 9.364389 | 0.105773 |
| **2.5_D** | 6.036595 | 5.880332 | 6.192252 | 0.079405 |
| **2.5_L** | 5.523959 | 5.317355 | 5.729247 | 0.104627 |
| **5_D** | 5.044999 | 4.810948 | 5.272399 | 0.106202 |
| **5_L** | 4.728456 | 4.549895 | 4.905853 | 0.0904 |
| **10_D** | 3.715673 | 3.480478 | 3.938372 | 0.110912 |
| **10_L** | 2.573887 | 2.273749 | 2.859186 | 0.069296 |

Treatment groups: Control\_None (10% sucrose + water), 0\_None (water), 2.5\_D (2.5% D-arabinose + water), 2.5\_L (2.5% L-arabinose + water), 5\_D (5% D-arabinose + water), 5\_L (5% L-arabinose + water), 10\_D (10% D-arabinose + water), and 10\_L (10% L-arabinose + water).

**Supp. Table 7:** All significantly different LT50 time points for various arabinose treatments in Ae. aegypti.

| **Contrast** | **Estimate** | **SE** | **t-ratio** | **p-value** |
| --- | --- | --- | --- | --- |
| **0 - 2.5_D** | 0.179858 | 0.031577 | 5.69589 | 2.59E-05 |
| **0 - Control** | 0.280725 | 0.030717 | 9.139213 | 0 |
| **2.5_D - 2.5_L** | -0.28109 | 0.028281 | -9.93889 | 0 |
| **2.5_D - 5_D** | -0.27349 | 0.025735 | -10.6274 | 0 |
| **2.5_D - 5_L** | -0.2537 | 0.028677 | -8.84659 | 0 |
| **2.5_L - Control** | 0.381953 | 0.027317 | 13.98203 | 0 |
| **5_D - Control** | 0.374359 | 0.024671 | 15.17394 | 0 |
| **5_L - Control** | 0.354564 | 0.027727 | 12.78757 | 0 |
| **10_D - 2.5_D** | 0.291696 | 0.031978 | 9.121856 | 0 |
| **10_D - Control** | 0.392563 | 0.031129 | 12.61102 | 0 |
| **10_L - 2.5_D** | 0.258692 | 0.030019 | 8.617719 | 0 |
| **10_L - Control** | 0.359559 | 0.029112 | 12.35077 | 0 |

Treatment groups: Control\_None (10% sucrose + water), 0\_None (water), 2.5\_D (2.5% D-arabinose + water), 2.5\_L (2.5% L-arabinose + water), 5\_D (5% D-arabinose + water), 5\_L (5% L-arabinose + water), 10\_D (10% D-arabinose + water), and 10\_L (10% L-arabinose + water).

**Supp. Table 8:** All significantly different LT50 timepoints for various arabinose treatments in Cx. pipiens.

| **Contrast** | **Estimate** | **SE** | **t-ratio** | **p-value** |
| --- | --- | --- | --- | --- |
| **0 - Control** | 0.395481 | 0.027386 | 14.44107 | 0 |
| **10_D - Control** | 0.409151 | 0.027798 | 14.71864 | 0 |
| **10_L - Control** | 0.476708 | 0.030168 | 15.80155 | 0 |
| **2.5_D - Control** | 0.358587 | 0.023579 | 15.20787 | 0 |
| **2.5_L - Control** | 0.361448 | 0.028522 | 12.67271 | 0 |
| **5_D - Control** | 0.34666 | 0.024736 | 14.01448 | 0 |
| **5_L - Control** | 0.370957 | 0.026626 | 13.9322 | 0 |

Treatment groups: Control\_None (10% sucrose + water), 0\_None (water), 2.5\_D (2.5% D-arabinose + water), 2.5\_L (2.5% L-arabinose + water), 5\_D (5% D-arabinose + water), 5\_L (5% L-arabinose + water), 10\_D (10% D-arabinose + water), and 10\_L (10% L-arabinose + water).

**Supp. Table 9:**
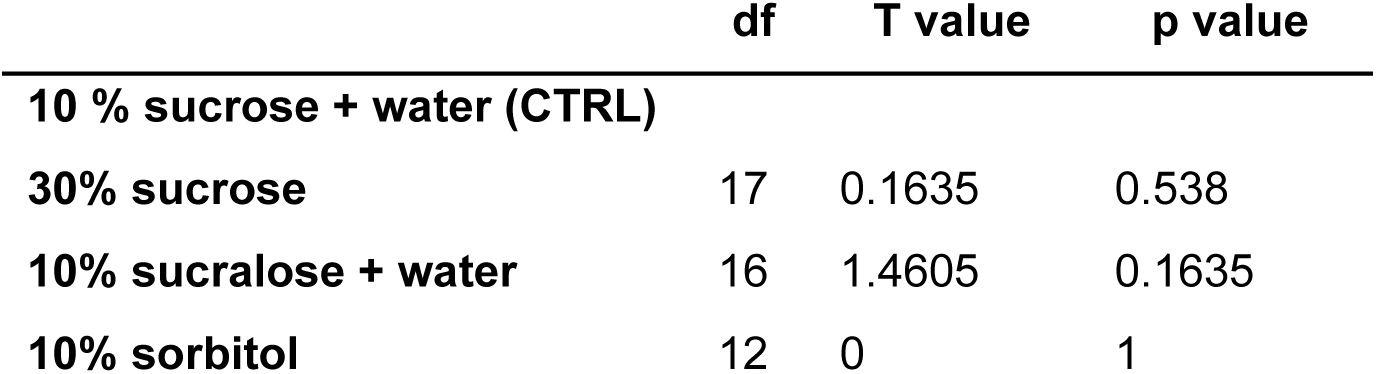
Host-seeking behaviors of *Ae. aegypti* after five days of sugar-diet treatments. No significance was found when the treatment groups were compared to the control group (CTRL, 10% sucrose + water) by t-test

